# The nuclear actin cytoskeleton supports DNA double-strand break repair via VCP-mediated extraction of the KU70/80 complex from damaged chromatin

**DOI:** 10.64898/2026.09.01.748302

**Authors:** Kristine Hauschulte, Jie Shi, Ronald P. Wong, Felizitas F. Stiehler, Sebastian Richenhagen, Aisha A. Kahloon, Lea-Marie Vogts, Markus S. Schraft, Tina Strauch, Stephanie Nick, Ivan Mikicic, Marton Gelleri, Fridolin Kielisch, Vassilis Roukos, Krishnaraj Rajalingam, Petra Beli, Alexander Loewer, Helle D. Ulrich, Hans-Peter Wollscheid

**Author notes:** Equal contribution (Kristine Hauschulte, Jie Shi). Lead contact & correspondence and.

## Abstract

Double-strand breaks (DSBs) are critical lesions in genomic DNA, and their accurate repair is essential for maintaining genome stability. The nuclear actin cytoskeleton has been implicated in homology-directed repair (HDR) of DSBs. However, the underlying mechanism remains poorly understood. Here, we report that Myosin VI (Myo6), an actin-based motor protein, cooperates with F-actin in end resection and DSB mobilization. Our findings reveal that Myo6 directly interacts with both KU70 and the ubiquitin-dependent segregase VCP to facilitate the extraction of the KU70/80 complex from chromatin. This process is supported by F-actin, revealing an interplay between nuclear actin dynamics and the DSB repair machinery. By elucidating the function of Myo6 and its direct interactions with key repair factors, our study provides mechanistic insight into how repair mechanisms rely on nuclear actin to safeguard genome integrity.

## Introduction

Filamentous actin (F-actin), traditionally recognized for its cytoplasmic roles, has emerged as a pivotal player within the nuclear environment, orchestrating chromatin dynamics and genome stability^1–3^. Nuclear F-actin assembly is induced by various DNA-damaging agents, suggesting an involvement in multiple aspects of the DNA repair process^4^. Although mechanistic insight is still missing, recent studies underscore a significant role of nuclear F-actin in the homology-directed repair (HDR) of DNA double-strand breaks (DSBs)^1,5–7^. DSBs represent highly toxic lesions that are primarily repaired through two distinct pathways^8^. During the G1 phase of the cell cycle, DSBs are predominantly resolved via error-prone non-homologous end-joining (NHEJ), mediated by the DNA end-binding KU70/80 complex. In S and G2 phase, a subset of DSBs is channeled into resection-dependent repair pathways including HDR, which is more accurate due to its reliance on a homologous sequence as a repair template^9^. Removal of the KU70/80 complex from chromatin is a prerequisite for DNA end resection and is facilitated by the AAA-ATPase VCP^10^. In human cells, inhibition of nuclear actin polymerization interferes with two important steps of the HDR pathway, DNA end resection and DSB mobilization^1^. Moreover, nuclear F-actin coordinates three-dimensional chromatin organization and chromatin accessibility, thereby modulating the transcriptional response to genomic insults^11–13^.

Due to technical challenges in the visualization and manipulation of nuclear actin filaments, most studies have so far relied on targeting actin-binding proteins, particularly actin nucleation factors such as formins or the ARP2/3 complex. Interfering with their activity results in reduced nuclear F-actin assembly and compromised DSB repair in yeast and higher eukaryotes ^4–7,14,15^. In *Drosophila* and mice, the nuclear ARP2/3 complex promotes actin assembly to drive DSB relocation, whereas in yeast and human cells, ARP2/3 regulates DNA end resection and DNA mobility^1,5,7^.

In contrast to F-actin, the role of myosins in DSB repair is poorly understood. While *Drosophila* myosins of class I and V mediate DSB relocation, contributions of human myosins to DSB repair have not been reported^16^. Among the few myosin family members known to have nuclear roles, Myosin VI (Myo6) has been observed to localize to the nucleus in response to DNA damage and was reported to exhibit pro-survival functions under conditions of severe genomic insult^17–19^. However, the precise mechanisms underlying its nuclear involvement remain elusive, and our understanding of how Myo6 contributes to DNA repair is still undeveloped.

Here, we report a role of Myo6, a member of the myosin superfamily with unique pointed-end directionality^20^ harboring a pair of ubiquitin-binding domains^21^, in the HDR pathway. We demonstrate that Myo6 plays crucial roles in the same two steps that are controlled by F-actin: DNA end resection and DSB mobility. Through proteomic data analysis, we identify interactions of Myo6 with the KU70/80 complex and the AAA-ATPase VCP. Using laser micro-irradiation experiments, we demonstrate ubiquitin-dependent recruitment of Myo6 to sites of DNA damage. Increased retention of a motor-less mutant suggests a transport function of Myo6 away from the break site. Consistent with the observed defects in DNA end resection, we detect a strong impairment in the resolution of KU80 foci after DNA damage in Myo6-deficient cells as well as upon pharmacological inhibition of the ARP2/3 complex. In summary, our investigation into Myo6’s role in DSB repair reveals an actin-myosin-dependent molecular mechanism that facilitates long-range DNA end resection and HDR of DSBs by supporting VCP-dependent extraction of KU70/80 from damaged chromatin.

## Results

### Myo6 contributes to the homology-directed repair pathway of DNA double-strand breaks

In a mass spectrometry-based screen for interactors of the Myo6 ubiquitin-binding domain in Hela cells, we recently identified several DNA replication factors, revealing a contribution of Myo6 to replication fork protection^22^. In addition, we identified multiple DNA repair factors, some of them apparently unrelated to DNA replication. GO enrichment analysis for biological processes identified 22 proteins involved in homologous recombination (HR) and six proteins related to non-homologous end joining (NHEJ) (Fig. 1a,b). We therefore decided to investigate a direct contribution of Myo6 to DSB repair. To this end, we challenged Myo6-depleted or *knockout* A549 cells (KO) with the DSB-inducing agents Camptothecin (CPT) and Neocarcinostatin (NCS) and compared their survival to siCTRL-treated or *wildtype* A549 cells (WT). While Myo6-deficient cells formed an equal number of colonies compared to control cells under unchallenged conditions, they showed increased sensitivity towards CPT and NCS (Fig. 1 c,d), suggesting a critical role of Myo6 in DSB repair. To test contributions to HDR versus NHEJ, we employed a reporter-based assay (Traffic Light Reporter, TLR) (Fig. 1e, Supplementary Fig. 1)^23^. CtBP-interacting protein (CtIP), a factor involved in the initiation of DNA end resection during the early stages of HDR, served as a control^24^. As expected, depletion of CtIP by siRNA resulted in a strong decrease in HDR efficiency (Fig. 1f, Supplementary Fig. 1). Similarly, but to lesser extent, overall DSB repair (NHEJ and HDR) activity was impaired by an ARP2/3 inhibitor, CK666, confirming previous findings regarding the function of F-actin during HDR^1^. In line with a potential function in HDR-mediated DSB repair, we observed compromised HDR efficiency in cells treated with siRNAs against Myo6 (Fig. 1f, Supplementary Fig. 1). In summary, our data suggest a yet undescribed function of the actin-based motor Myo6 in promoting the homology-directed repair of DSBs.

**Fig. 1.**
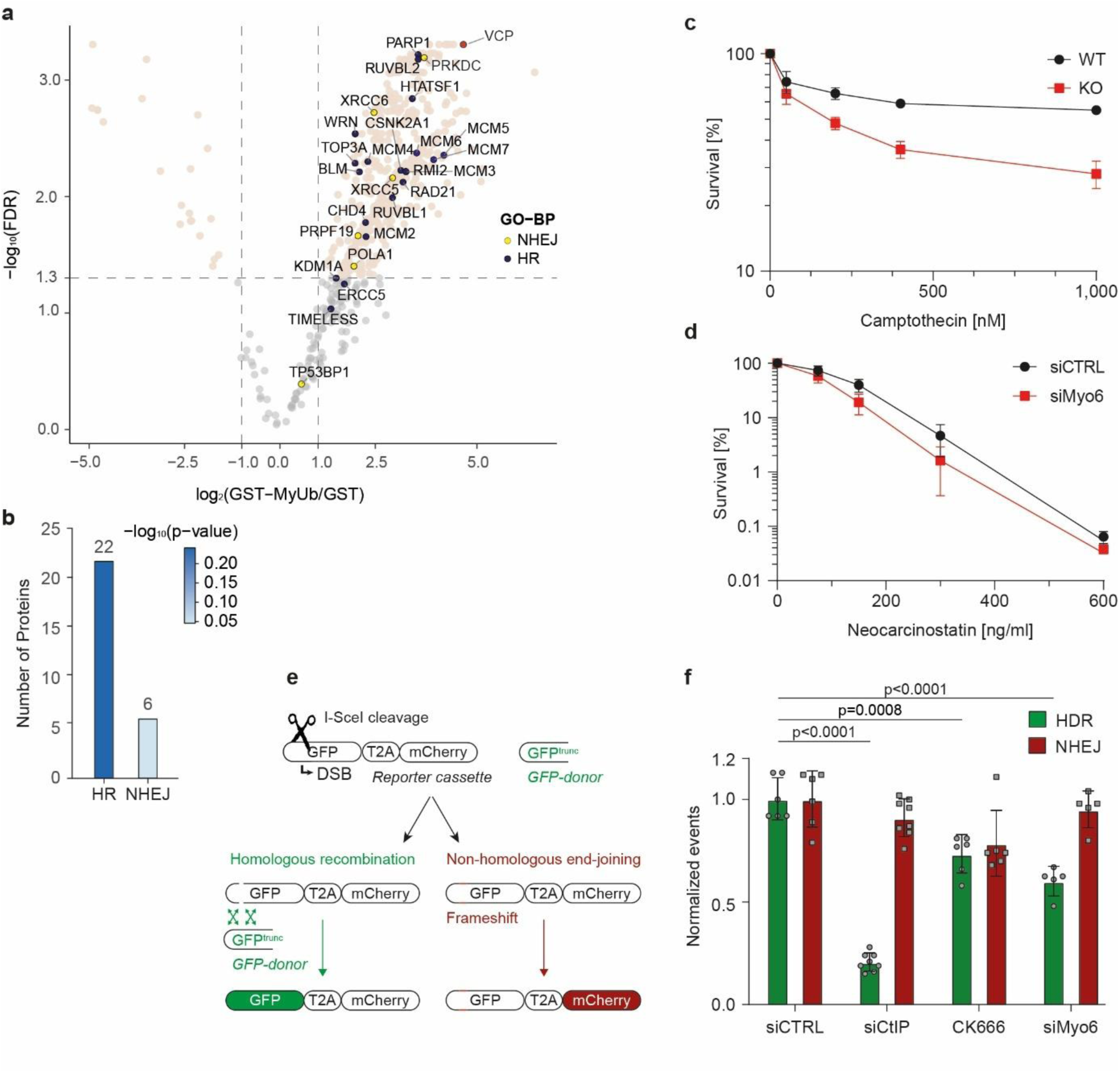
Myo6 is involved in the homology-directed DSB repair pathway. **a**, Myo6 interacts with DSB repair proteins. Volcano plot of protein groups identified as interactors of the Myo6 MyUb domain by SILAC-based mass spectrometry, reanalyzed from Shi et al.^22^. Mean log_2_ fold changes of all three replicates of GST-MyUb versus GST are plotted against the −log_10_(FDR). Significantly enriched proteins are shown in beige (fold change > 2, FDR < 0.05). Enriched interactors involved in GO biological processes (clusterProfiler) “homologous recombination” (HR) and “non-homologous end joining” (NHEJ) are highlighted and labeled. **b,** Bar plot summarizing GO biological process enrichment for proteins identified in the MyUb interactome shown in panel a. GO terms for “homologous recombination” (HR) and “non-homologous end joining” (NHEJ) were queried against the filtered interactome using clusterProfiler. The number of enriched proteins assigned to each pathway and −log_10_(p-value) are displayed. **c, d,** Myo6-depleted cells show increased sensitivity to DSB-inducing agents. (c) A549 *wildtype* (WT) and Myo6 *knockout* (KO) cells were treated with increasing concentrations of Camptothecin (CPT), as indicated. (d) A549 cells were transfected with the indicated siRNAs and exposed to increasing concentrations of Neocarcinostatin (NCS) 72 h after transfection. Viability was assessed by crystal violet staining. Mean values −/+ 95% confidence intervals from at least three independent experiments are shown. **e,** Schematic representation of the reporter assay used in panel **f** (modified from *Certo et al*.^23^). **f,** Depletion of Myo6 reduces HDR efficiency. U2OS cells harboring the Traffic Light reporter as illustrated in panel e were transfected with plasmids encoding the I-SceI nuclease and a GFP donor. Cells were either treated with 50 µM of the ARP2/3 inhibitor CK666 for 72 h or co-transfected with siRNAs against CtIP or Myo6 as indicated and analyzed by flow cytometry 72 h post-transfection. Repair efficiencies are shown relative to siCTRL-treated cells as individual data points with mean values (bars) and −/+ 95% confidence intervals. Significance levels were calculated using the two-tailed t test.

### Myo6 supports DNA end resection and DSB mobility

Nuclear F-actin supports the HDR pathway by ensuring efficient DNA end resection and facilitating DSB mobility required for break clustering^1^. To test if Myo6 engages in these processes as well, we measured DSB mobility using an established reporter system based on U2OS cells expressing a fusion of RAD52 with the red-fluorescent protein mCherry^25^. We challenged reporter cells with ionizing radiation to induce DSBs, allowed 20 h for repair pathway activation and then followed the movement of RAD52 foci for 80 min. We first quantified DSB mobility using mean-square displacement (MSD) analyses in the presence or absence of the ARP2/3 inhibitor CK666 (Fig. 2a). In line with previously published data^1^, we observed reduced DSB mobility under these conditions, confirming the involvement of F-actin in mobilizing DSBs (Fig. 2a). Interestingly, we observed an even stronger reduction in DSB mobility when downregulating Myo6 expression using siRNA, supporting our hypothesis that Myo6 cooperates with nuclear F-actin to promote resection-dependent repair (Fig. 2b).

**Fig. 2.**
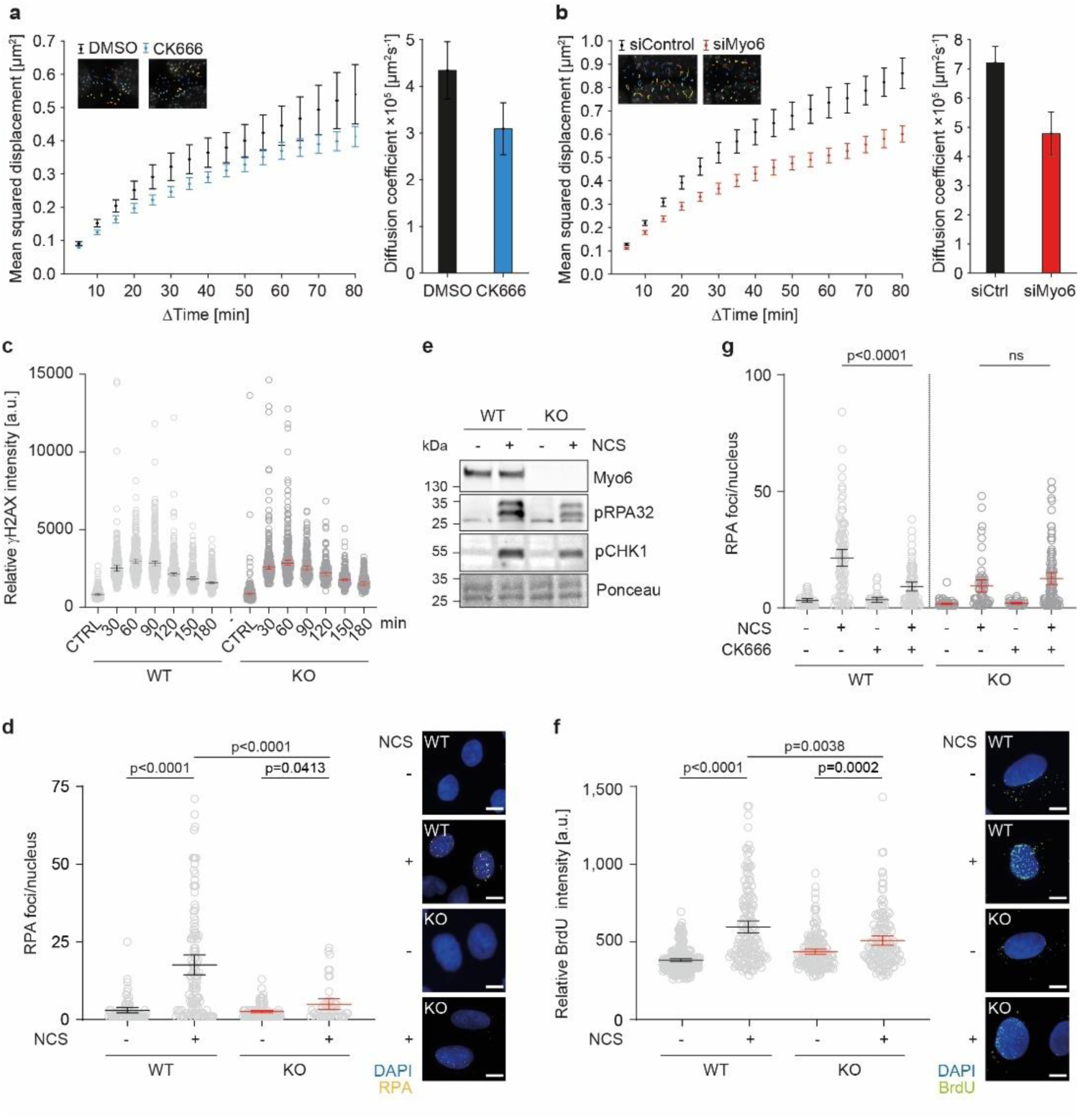
Myo6 regulates DNA end resection and DSB mobility. **a, b**, DSB mobility is reduced upon perturbation of Myo6 and F-actin. U2OS cells expressing Rad52-mCherry were irradiated with 5 Gy X-ray, Rad52 foci were followed by live-cell microscopy for 80 min at 5 min intervals 20 h post-damage. Mean squared displacement (MSD) is shown as weighted mean; error bars indicate weighted standard deviation. Diffusion coefficients were calculated from linear fits to the first 30% of the weighted mean MSD curve; error bars indicate 95% confidence intervals. Insets show images of representative nuclei including traces of Rad52 foci. Cells were either treated with DMSO and the ARP2/3 inhibitor CK666 (a) or transfected with siRNAs against Myo6 (b). n = 704 from 20 nuclei (DMSO), 776 from 20 nuclei (CK666), 1118 from 36 nuclei (siCTRL) 1162 from 37 nuclei (siMyo6). Data were pooled from 2-3 independent replicates. **c,** Myo6 does not affect H2AX phosphorylation. A549 WT and Myo6 KO cells were treated with 1 µg/ml NCS for 1 h, and γH2AX intensities were followed by immunofluorescence microscopy over 3 h in 30 min intervals. Signal intensities were quantified using Image J and shown as dot plots with mean values −/+ 95% confidence intervals. **d,** Myo6 promotes RPA foci formation upon DSB induction. A549 WT and Myo6 KO cells were treated with 1 µg/ml NCS for 1 h followed by immunofluorescence microscopy against RPA32. **e,** Myo6 promotes RPA32 phosphorylation upon DSB induction. A549 WT and Myo6 KO cells were treated with 1 µg/ml NCS for 2 h. Total cell lysates were analyzed by western blotting and Ponceau S staining. **f,** Myo6 promotes ssDNA formation upon DSB induction. A549 WT and Myo6 KO cells were labeled with BrdU for 24 h, treated with 1 µg/ml NCS for 1 h and analyzed by immunofluorescence microscopy under non-denaturing conditions using a BrdU-specific antibody. **g,** Loss of Myo6 is epistatic with F-actin inhibition in RPA foci formation. A549 WT and Myo6 KO cells were treated with either DMSO or 100 µM ARP2/3 inhibitor CK666 and analyzed as in panel **e**. For **d+f,** Representative images are shown. Scale bars = 10 µm. For **c-g,** A representative experiment from at least three independent replicates is shown. For **d, f, g,** Signal intensities were quantified using Image J and shown as dot plots with mean values −/+ 95% confidence intervals. Significance levels were calculated using the Mann-Whitney test (ns: not significant).

Next, we examined the effect of Myo6 on DNA end resection by monitoring a series of characteristic steps in the process. Phosphorylation of the histone variant H2AX marks the activation of damage signaling upstream of resection^26^. Here, as expected, Myo6-deficient cells did not show a defect in DSB recognition (Fig. 2c). To study resection-dependent formation of single-stranded DNA (ssDNA), we used various approaches, starting with microscopy-based analyses of the heterotrimeric Replication Protein A (RPA), the major ssDNA-binding protein^27^. Myo6 deletion resulted in a reduced RPA foci count compared to WT cells, supporting a potential function of Myo6 in DNA end resection (Fig. 2d, Supplementary Fig. 2a). Upon chromatin-binding, RPA undergoes PI3 kinase-mediated phosphorylation of defined serine and threonine residues within the N-terminus of the RPA32 subunit, indicating activation of the DNA damage checkpoint^27^. While control siRNA-treated cells showed a robust induction of RPA phosphorylation after exposure to CPT or NCS, Myo6 KO cells exhibited a weaker response (Fig. 2e, Supplementary Fig. 2b).

To validate our findings by means of an RPA-independent assay to detect ssDNA, we visualized BrdU-labeled DNA by immunofluorescence under native conditions^28^. This method relies on the inability of the BrdU-specific antibody to bind its antigen in the context of dsDNA and therefore specifically recognizes ssDNA under non-denaturing conditions. Following a 24-hour labeling period, we observed diminished BrdU signals in Myo6 KO cells after DNA damage induction compared to control cells (Fig. 2f, Supplementary Fig. 2c). To further assess the downstream effects of Myo6 depletion on DNA repair processes, we also monitored RAD51 foci formation, an event downstream of DNA end resection. As expected, a reduction in chromatin-bound RAD51 was observed in Myo6 KO cells after DSB induction with NCS (Supplementary Fig. 2d). These results collectively demonstrate that Myo6-deficient cells exhibit impaired DNA end resection, leading to attenuated damage signaling and limited downstream processing.

To explore whether the function of Myo6 during DNA end resection depends on its cooperation with F-actin, we tested whether Myo6 depletion and simultaneous inhibition of ARP2/3-dependent actin polymerization would result in additive or epistatic effects on the formation of damage-induced RPA32 foci. Single inactivation of either F-actin or Myo6 caused a prominent decline in the number of RPA32 foci, and combining the two conditions did not lead to any further decrease (Fig. 2g). This suggests a cooperation of Myo6 with F-actin during DNA end resection. In summary, we established Myo6 as a regulator of DSB mobility and end resection, suggesting a function for this motor protein in the HDR pathway, possibly in transporting cargo on the short, transient nuclear actin polymers that were previously reported to accompany the processing of DSBs^1^.

### Myo6 localizes to DNA damage sites in a ubiquitin-dependent manner

Formation and function of actin filaments inside the nucleus are still controversially discussed^2^. Therefore, we asked whether the contribution of Myo6 to DNA end resection and DSB mobility is direct or mediated indirectly via its predominant cytoplasmic pool. Due to its low nuclear abundance, we failed to detect co-localization of endogenous Myo6 with classical DSB markers, including γH2AX and 53BP1, using immunofluorescence microscopy. Hence, we decided to generate larger areas of DNA damage via UV laser-based micro-irradiation to facilitate the detection of repair factors recruited to the lesions. We performed time-lapse imaging of cells overexpressing GFP-Myo6 and monitored the dynamics of laser-induced Myo6 recruitment to the damaged area. In undamaged cells, GFP-Myo6 was largely cytoplasmic without a detectable nuclear signal, but in agreement with previously published data^17^, laser-induced DSB formation resulted in a rapid overall relocalization to the nucleus (Fig. 3a). Moreover, we detected a robust enrichment of Myo6 at the damaged chromatin area. To determine whether the motor activity of Myo6 is required for this recruitment, we tested a truncated, motor-deficient mutant (GFP-Myo6Δ1-841, here named as “GFP-Tail”) under the same conditions. UV laser micro-irradiation induced recruitment with comparable kinetics (Supplementary Fig. 3) but enhanced retention of the truncated protein at the site of damage compared to its full-length counterpart (Fig. 3b and Supplementary Fig. 3). This unexpected result suggests that Myo6’s motor function is not important for recruitment or retention at the sites of DNA damage, but rather for its release.

**Fig. 3.**
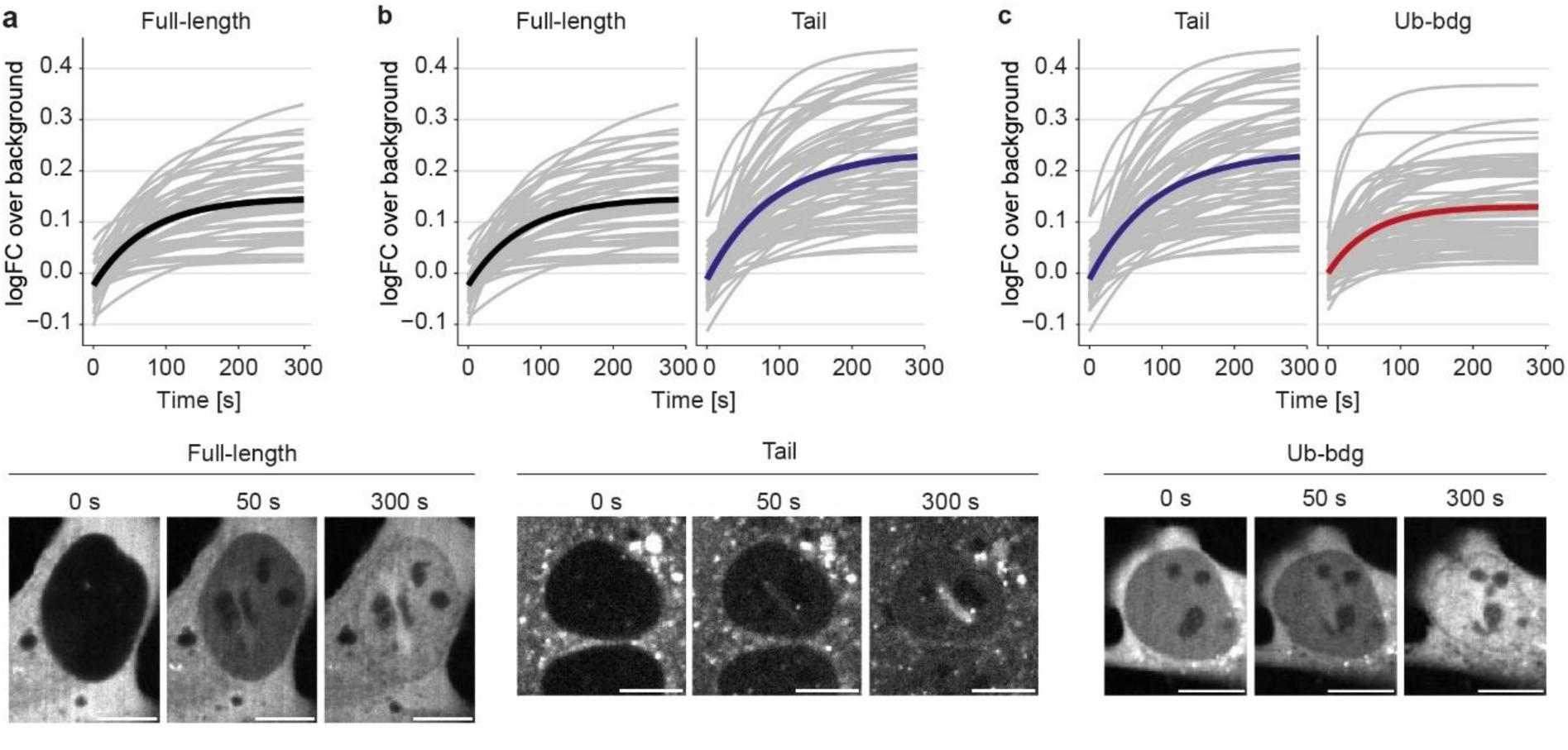
Localization of Myo6 to DSBs depends on ubiquitin binding. **a**, Myo6 is recruited to sites of UV laser-induced damage. U2OS cells, transiently transfected with full-length GFP-Myo6, were subjected to UV laser micro-irradiation. Enrichment of GFP-Myo6 at the damaged region was followed by time-lapse microscopy over 300 s in 5 s intervals. **b,** A defect in the motor domain enhances retention of Myo6 at sites of DNA damage. U2OS cells were transiently transfected with a motor-deficient GFP-Tail construct (Tail) and subjected to UV laser micro-irradiation as in panel **a**. WT values from panel a are shown for comparison. **c,** A defect in the ubiquitin-binding domain interferes with recruitment of Myo6 to sites of DNA damage. U2OS cells, transiently transfected with GFP-Myo6 construct deficient in ubiquitin binding (Ub-bdg, A1013G, I1104A) and subjected to UV laser micro-irradiation as in panel **a**. Values for the Tail domain from panel b are shown for comparison. For **a-c,** GFP signals were analyzed using Image J. The relative increase of the GFP signal at the stripe over the GFP signal in the undamaged chromatin region was plotted as mean (highlighted curve) from all single measurements (light grey curves) from three independent replicates. Representative images for 0, 50, and 300 s are shown. Scale bars = 10 µm.

The DNA damage response (DDR) involves a series of ubiquitylation events, mediated by – among others – the ubiquitin protein ligases (E3s) RNF8 and RNF168 and resulting in highly ubiquitin-decorated chromatin at and around the break site^29^. To study the importance of the tandem ubiquitin-binding domains of Myo6 (MIU and MyUb)^30^ for its recruitment to DSBs, we introduced two point mutations, A1013G (MIU domain) and I1104A (MyUb domain)^21,31^, into the GFP-Tail construct to interfere with its ubiquitin-binding activity. These mutations strongly reduced recruitment to damaged chromatin (Fig. 3c).

In summary, using UV laser micro-irradiation, we observed physical recruitment of Myo6 to DSB sites in a manner that is independent of its motor domain but is supported by Myo6’s ubiquitin-binding activity. The notion that a motor-deficient Myo6 mutant exhibited enhanced retention at the damage implies an actin-dependent motor activity in a transport-related function directed away from the DSB.

### Myo6 directly interacts with VCP and the KU70/80 complex

Myo6-dependent transport of chromatin-associated factors away from the break site could in principle facilitate the removal of negative regulators of HDR or the turnover of repair enzymes to ensure the correct sequence of temporally orchestrated events. Among potential candidates whose removal would promote HDR are the NHEJ factors, KU70 (XRCC6) and KU80 (XRCC5). The KU70/80 complex recognizes free DNA ends of DSBs with high affinity and is an essential component of the NHEJ pathway. At the same time, it counteracts HDR by posing a physical barrier for helicases and nucleases driving DNA end resection^32^. Removal of the KU70/80 complex from chromatin to allow efficient HDR is in turn promoted by the AAA-ATPase VCP, a ubiquitin-dependent unfoldase that generally recognizes its substrates via adaptor proteins^30^(Fig. 4a). Accordingly, the KU proteins undergo polyubiquitylation prior to their extraction. Remarkably, KU70, KU80, and VCP were all identified as Myo6 interactors in our proteomic dataset with high confidence (Fig. 1a). Considering the end resection and HDR defect of Myo6-deficient cells, we asked whether Myo6 exerts its function in HDR via the KU70/80 complex and/or VCP.

**Fig. 4.**
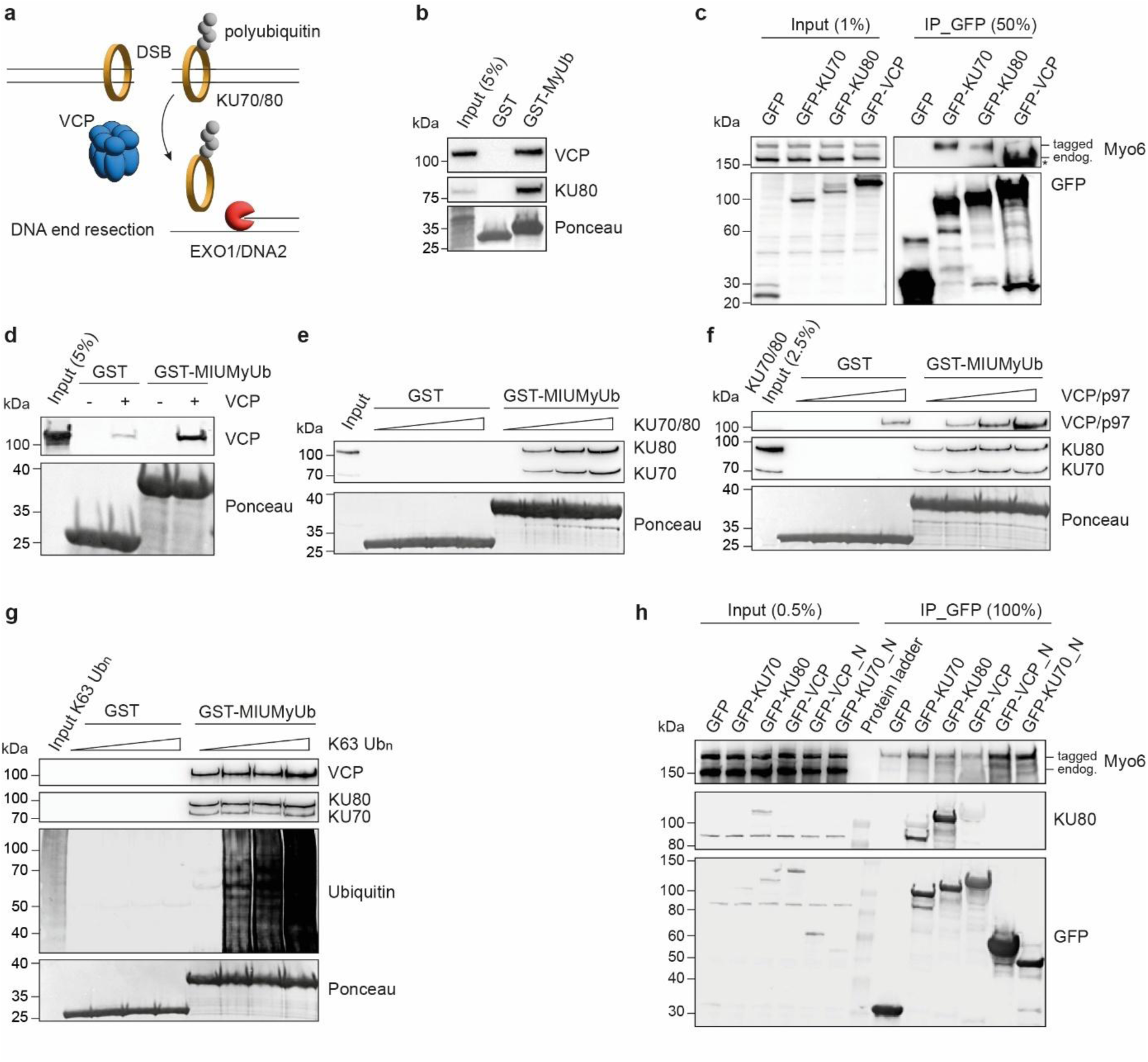
Myo6 directly interacts with the KU70/80 complex and VCP. **a**, Schematic representation showing how VCP-mediated extraction of the KU70/80 complex from chromatin promotes efficient long-range resection. **b,** Validation of the interaction of Myo6 with KU80 and VCP by pulldown assays from total cell lysates using recombinant GST-MyUb, followed by western blotting and Ponceau S staining. GST served as a negative control. **c,** Myo6 binds the KU70/80 complex in cellular lysates. HEK293T cells were co-transfected with DamID-FLAG-tagged Myo6 and the indicated GFP-fusion constructs and were subjected to immunoprecipitations (IPs) against GFP, followed by western blotting. The asterisk marks cross-reactivity of the anti-Myo6 antibody with GFP-VCP. GFP served as a negative control. **d,** Myo6 directly interacts with VCP. A pulldown assay was performed with recombinant GST or GST-Myo6 UBDs (GST-MIUMyUb) as a bait and recombinant full-length VCP. Proteins were visualized by western blotting and Ponceau S staining. **e,** Myo6 directly interacts with the KU70/80 complex. A pulldown assay was performed with GST or GST-Myo6 UBDs (GST-MIUMyUb) as a bait and a 4-fold concentration range of recombinant KU70/80 complex. Input represents 20% of the lowest concentration used. Proteins were visualized by western blotting and Ponceau S staining. **f,** Myo6 simultaneously binds to KU70/80 and VCP. A GST-pulldown assay was performed with GST-Myo6 UBDs (GST-MIUMyUb) and recombinant KU70/80 complex and a 4-fold concentration range of full-length VCP. Proteins were visualized by western blotting and Ponceau S staining. **g,** Myo6 simultaneously binds to KU70/80, VCP, and ubiquitin. A GST-pulldown assay was performed with GST-Myo6 UBDs (GST-MIUMyUb) and recombinant full-length VCP, KU70/80 complex and a 4-fold concentration range of K63-linked polyubiquitin chains. Input represents 30% of the lowest concentration used. Proteins were visualized by western blotting and Ponceau S staining. **h,** Myo6 binds to the N-terminal regions of KU70 and VCP. HEK293T cells were transfected with DamID-FLAG-tagged Myo6 together with the indicated GFP fusion constructs and subjected to immunoprecipitations (IP) against GFP, followed by western blotting. **b-h,** Results were confirmed by at least two independent experiments.

To validate interactions of Myo6 with the KU70/80 complex and VCP, we first performed pulldown assays with the same GST-MyUb construct used in our previous SILAC screen with cellular extracts as a protein source. All tested interactions were confirmed by western blotting using target-specific antibodies (Fig. 4b). Next, we sought to confirm these interactions with full-length Myo6 using co-immunoprecipitation experiments. Isolation of GFP-tagged proteins with GFP-trap beads showed a prominent interaction between Myo6 and the KU70/80 complex, whereas the Myo6-VCP interaction was difficult to assess due to a strong cross-reactivity of the anti-Myo6 antibody (Fig. 4c). To assess whether the observed interactions were direct, we purified all proteins from bacteria and performed *in vitro* pulldown experiments using these recombinant proteins. For Myo6, we used a construct containing both UBDs (GST-MIUMyUb). Both VCP and the KU70/80 complex bound directly to this fragment (Fig. 4d, e). Moreover, the pre-assembled KU70/80-Myo6 interaction was not impeded by the addition of increasing amounts of VCP, indicating that concomitant binding of the KU70/80 complex and VCP to Myo6 is possible (Fig. 4f).

To assess whether Myo6 can accommodate binding to ubiquitin in addition to KU70/80 and VCP, we titrated pre-assembled ubiquitin chains into the Myo6-KU70/80-VCP binding assays. The notion that addition of ubiquitin chains did not displace VCP or KU70/80 from the Myo6 fragment (Fig. 4g) is consistent with Myo6 binding to ubiquitylated KU70/80 complex and VCP simultaneously, potentially to facilitate efficient extraction of the KU70/80 complex from damaged chromatin.

We used AlphaFold 3^33^ to identify potential binding surfaces on KU70/80 and VCP for their interaction with Myo6. In this manner, interactions of Myo6’s UBDs with the N-termini of KU70 and VCP, but not with KU80, were predicted with high confidence (Supplementary Figs. 4, 5). To substantiate these predictions, we performed immunoprecipitation experiments using the respective full-length as well as truncated proteins. Accordingly, we observed a weaker interaction of Myo6 with GFP-KU80 compared to GFP-KU70, suggesting a KU70-dominated binding mode. In line with the AlphaFold predictions, we found that an N-terminal KU70 fragment (aa1-249) still bound to Myo6, while its interaction with KU80 was lost (Fig. 4h, Supplementary Fig. 4). Furthermore, we scored a prominent increase in the interaction between Myo6 and the N-terminal fragment of VCP compared to the full-length protein (Fig. 4h), again consistent with the AlphaFold prediction (Supplementary Fig. 5). Collectively, our data reveal direct interactions between Myo6, KU70 and VCP, suggesting a potential function of Myo6 in the VCP-mediated extraction of the KU70/80 complex from chromatin during the initial steps of HDR to promote efficient long-range DNA end resection.

### Myo6 assists in VCP-mediated removal of KU70/80 complex from damaged chromatin

Involvement of Myo6 in the VCP-mediated removal of the KU70/80 complex from damaged chromatin should result in enhanced retention of the KU proteins on chromatin upon inactivation of Myo6, analogous to what was previously shown for VCP inhibition^10^. To test this prediction, we subjected A549 WT and Myo6 KO cells to ionizing radiation and followed KU80 foci formation and resolution over time using immunofluorescence. In WT cells, we detected KU80 foci within one hour after irradiation and a gradual resolution over time, reflecting repair of the breaks (Fig. 5a). VCP inhibition afforded no changes in the formation of KU80 foci; however, in line with the role of VCP in the removal of the KU70/80 complex, foci resolution was compromised at later time points. Notably, a similar persistence of KU80 on chromatin was observable in Myo6 KO cells, supporting our hypothesis that Myo6 contributes to removal of the KU70/80 complex (Fig. 5a, c).

**Fig. 5.**
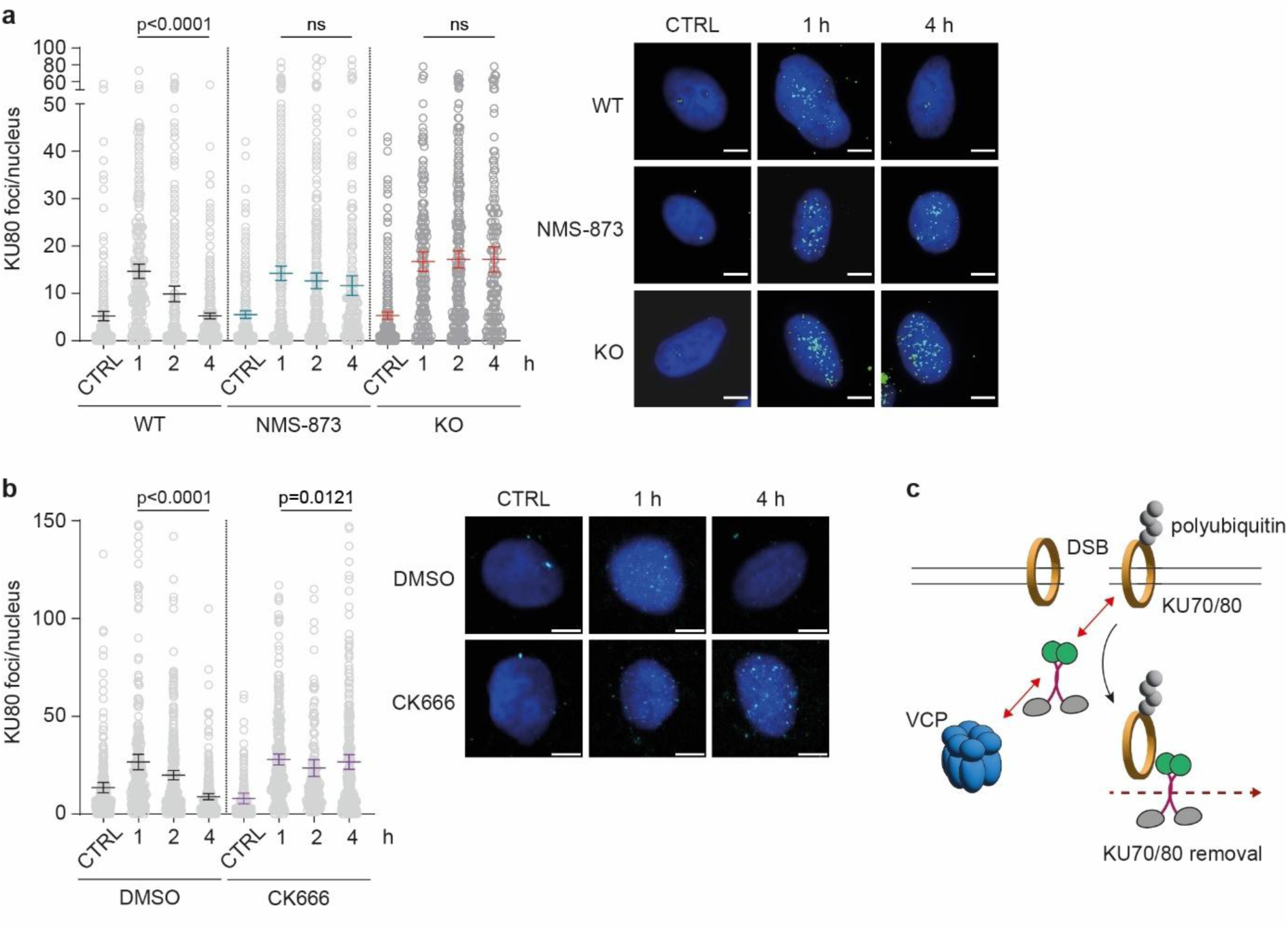
The actin cytoskeleton promotes removal of KU70/80 from chromatin. **a**, Myo6 and VCP are required for efficient resolution of KU80 foci after DNA damage. A549 WT and Myo6 KO cells were exposed to 10 Gy X-irradiation. KU80 chromatin retention was followed by immunofluorescence microscopy at the indicated time points after irradiation. Where indicated, 5 µM VCP inhibitor NMS-873 was added to A549 WT cells 1 h prior to irradiation. **b,** F-actin contributes to the resolution of KU80 foci after DNA damage. A549 cells were exposed to 10 Gy X-irradiation. KU80 chromatin retention was followed by immunofluorescence microscopy at the indicated points after irradiation. Where indicated, 100 µM ARP2/3 inhibitor CK666 or DMSO was added to A549 WT cells 1 h prior to irradiation. **c,** Model illustrating how Myo6 and F-actin could facilitate VCP-mediated removal of KU70/80 complexes from damaged chromatin. Myo6 mediates the interaction (red arrows) between VCP and KU70/80. After successful chromatin extraction, Myo6 removes the KU70/80 complex from the damaged area to avoid re-binding. **a, b,** Representative images are shown on the right. Image data were analyzed using Image J and shown as dot plots with mean values −/+ 95% confidence intervals. Significance levels were calculated using the Mann-Whitney test (ns: not significant). Scale bars = 10 µm.

Assuming an F-actin-dependent transport activity of Myo6 in this process, we predicted that inhibition of the actin polymerization factor ARP2/3 would likewise interfere with the extraction of the KU70/80 complex. Notably, an interaction between F-actin and the KU70/80 complex has been described previously^14^. As expected, the ARP2/3 inhibitor, CK666, also prevented K80 foci resolution (Fig. 5b). Taken together, our observations suggest a cooperation of nuclear F-actin with Myo6 during VCP-mediated extraction of the KU70/80 complex from DSBs to allow efficient resection and HDR (Fig. 5c).

## Discussion

DNA double-strand breaks (DSBs) pose a significant threat to genome stability, necessitating precise repair mechanisms. By implicating a human actin-based motor protein in critical steps of resection-dependent pathways, our study provides a mechanistic link between the actin cytoskeleton and the process of DSB repair.

Here, we demonstrate that Myo6 promotes DNA end resection and DSB mobility, aligning with the established roles of nuclear F-actin in HDR. The recruitment of Myo6 to the nucleus and its accumulation at damaged chromatin suggest a direct involvement at the site of the break. Moreover, we provide evidence for physical interactions between Myo6, KU70, and VCP implying a mechanism where Myo6 cooperates with VCP in the removal of the KU70/80 complex in preparation for end resection. This is supported by the accumulation of chromatin-bound KU80 in Myo6 KO cells, mirroring effects seen with VCP inhibition.

Regarding the molecular details of how the nuclear actin cytoskeleton facilitates VCP’s action, several possibilities emerge. In one potential scenario, Myo6 could exert a “pulling” force or serve as an anchoring tether for VCP during the extraction of the KU70/80 complex from chromatin. However, this hypothesis seems inconsistent with the notion that VCP is known to act by pulling its substrate through its internal core via its ATPase activity, rendering external pulling forces redundant. Furthermore, direct recognition of KU70 by Myo6 would not be essential in such a scenario. Our findings suggest an alternative model, where Myo6 would serve as an adaptor protein mediating interactions between VCP and the ubiquitylated KU70/80 complex, thereby promoting removal of the latter (Fig. 5c). This hypothesis is supported by Myo6’s ability to simultaneously interact with KU70, ubiquitin, and VCP. Previous studies by van den Boom et al. demonstrated that KU80 undergoes polyubiquitination in response to DNA damage and is recognized by classical VCP adaptors, UFD1/NPL4^10^. Consequently, depletion of these adaptors results in the accumulation of nuclear KU80 foci following DNA damage. In our proposed model, Myo6 may facilitate VCP’s recruitment to DSBs or enhance its ability to recognize the ubiquitylated KU70/80 complex as a substrate. In cooperation with F-actin, Myo6 might even play a dual role by not only aiding in the removal of the KU70/80 complex but also preventing its re-binding to DNA ends after extraction. This would ensure efficient and accurate repair during homology-directed repair processes.

In its simplest mode, this scenario would imply a direct recognition of the ubiquitylated KU70/80 complex by means of Myo6’s ubiquitin-binding domains. However, the preference of these domains for K63-linked ubiquitin chains argues against this scenario. We therefore favor a model where recruitment of Myo6 to DSBs relies on the complex chromatin-associated ubiquitin network initiated by the ubiquitin ligases RNF8 and RNF168^29^.

Our SILAC-based Myo6 interactome has served as a valuable resource not only for this study but also for uncovering a Myo6-supported, ubiquitin-dependent mechanism of replication fork protection^22^. The identification of other recombination and repair proteins, including BLM and WRN helicases, as well as chromatin remodeling factors such as RUVBL1/2 or CHD4 (Fig. 1a), raises the possibility that Myo6 and F-actin may have additional roles in homologous recombination or even other genome maintenance pathways. Once KU proteins are effectively cleared from DNA break ends, pre-existing actin filaments and their associated factors, including Myo6, may actively contribute to the resection process. Notably, in protecting stalled replication forks, Myo6 cooperates with Werner interacting protein 1 (WRNIP1) to limit the unscheduled activity of DNA2, a key nuclease involved in long-range resection^22^. Considering the unique directionality of Myo6, it is likely that other, plus-end directed motors also engage in the HDR pathway of human cells to promote DSB repair, potentially counteracting the effects of Myo6. Particularly interesting candidates are myosin type I and V, which transport heterochromatic DSBs to the nuclear periphery to promote error-free repair in *Drosophila*^5^. Future studies are required to determine the full extent to which the nuclear actin cytoskeleton impinges on chromatin dynamics during DNA replication and repair.

In summary, we have established Myo6 as an actin-binding protein involved in DNA double-strand break (DSB) mobility and DNA end resection. Through the study of Myo6, we have elucidated fundamental principles underlying how the nuclear actin cytoskeleton contributes to resection-dependent repair of DSBs.

## Methods

### Cell lines, cultivation and treatments

A549 *wildtype* and Myo6 *knockout* cells, U2OS, HeLa and HEK293T cells were maintained in DMEM containing 10% fetal bovine serum, L-glutamine (2 mM), penicillin (100 U/ml), and streptomycin (100 µg/ml) (Thermo Fisher Scientific). U2OS Flp-In T-REx cell lines and U2OS cells containing a stably integrated Traffic Light reporter cassette were maintained in DMEM containing 10% fetal bovine serum, L-glutamine, penicillin, streptomycin and blasticidin (5 µg/ml) (Invivogen). All cell lines were cultured in humidified incubators at 37°C with 5% CO_2_. Treatments were performed with neocarcinostatin (NCS), camptothecin (CPT), CK666 and NMS-873 at the indicated concentrations.

### Transfections

For overexpression purposes, HEK293T were transfected using polyethyleneimine (PEI) (Polysciences). All other cell types were transfected using Fugene HD (Promega) or Lipofectamine 2000 (Life technologies) according to the manufactureŕs instructions. All expression constructs used in this study are listed in Supplementary Table 1.

For knockdowns, cells were transfected with siRNAs using Lipofectamine RNAiMAX (Life Technologies) according to the manufactureŕs instructions at a final RNA concentration of 20 nM for 24-72 h, as indicated. Knockdown of Myo6 was achieved with a pool of 4 different siRNAs (Hs_MYO6_5 FlexiTube siRNA, Hs_MYO6_7 FlexiTube siRNA, Hs_MYO6_8 FlexiTube siRNA and Hs_MYO6_10 FlexiTube, Qiagen). CtIP knockdown was performed using a predesigned siRNA (Supplementary Table 3).

### Generation of plasmids

Fragments were inserted via restriction/ligation cloning following PCR amplification with specific oligonucleotides, listed in Supplementary Table 4. Detailed information about individual constructs will be provided upon request.

### Site-directed mutagenesis

Site-directed mutagenesis was performed using Pfu Turbo DNA Polymerase (Agilent). The amplification product was digested with DpnI (New England Biolabs), *E. coli* TOP10 cells were transformed with the construct followed by sequence verification. Oligonucleotides for mutagenesis are listed in Supplementary Table 4.

### Immunofluorescence

For immunofluorescence analysis, cells were fixed with 4% paraformaldehyde (PFA, Merck) for 10 min, permeabilized for 5 min at room temperature with 0.1% Triton X-100 and incubated for 1 h in PBS/3% BSA. Subsequently, cells were incubated with primary antibodies for 1 h (α-Myo6 in a 1:400 dilution), followed by 3 × 5 min washing steps with PBS/0.1% Triton X-100 and incubation with secondary antibodies for 30 min at room temperature. Coverslips were mounted with ProLong™ Diamond Antifade Mountant (Thermo Fisher Scientific). Images were acquired with a Leica AF-7000 widefield microscope and quantified using custom macros in Fiji/ImageJ.

For RPA and Rad51 staining, cells were washed with PBS and pre-extracted with 0.3 % IGEPAL for 5 min on ice, washed in PBS twice and fixed in 4%. For BrdU staining, cells were washed with PBS and pre extracted with CSK A buffer (100 mM PIPES pH 6.8; 1 mM EGTA; 100 mM NaCl; 300 mM sucrose; 0,5% Triton) for 5 min on ice, washed in PBS twice and fixed in 4%. For KU80 staining, cells were washed with PBS and pre-extracted with CSK B buffer (10 mM PIPES, 100 mM NaCl, 300 mM sucrose, 3 mM MgCl2, 1% Triton-X, 0.3 mg/ml RNase A) for 5 min at RT, washed in PBS twice and fixed in 4% PFA.

### Immunoprecipitation of GFP-tagged proteins

HEK293T cells were PEI-transfected with the respective plasmid (Supplementary Table 1) for 24 h, followed by lysis in JS buffer (100 mM HEPES pH 7.5, 50 mM NaCl, 5% glycerol, 1% Triton X-100, 2 mM MgCl_2_, 5 mM EGTA, 1 mM DTT) supplemented with protease inhibitor cocktail (SIGMAFAST) and Benzonase®. Cell lysates were cleared by centrifugation for 30 min at 4°C and incubated with GFP-trap magnetic agarose beads (Chromotek) for 1 h at 4°C. After 3 washes with JS buffer, beads were boiled for 10 min in NuPAGE^®^ LDS Sample Buffer and subjected to western blotting.

### Immunoblotting

Samples were separated via SDS-PAGE and transferred to nitrocellulose membranes using the Trans-Blot Turbo® system (Bio Rad). Membranes were blocked for 1 h at room temperature in 5% milk/PBS/0.1% TWEEN-20 and incubated with primary antibodies (1:1000 dilution in PBS/0.1% TWEEN-20/1% BSA) either for 1 h at room temperature or overnight at 4°C. Afterwards, membranes were washed with PBS/0.1% TWEEN-20 and incubated with secondary antibodies for 1 h at room temperature. Detection was performed by enhanced chemiluminescence using a Fusion FX (Vilber Lourmat) instrument after incubation with HRP-coupled secondary antibodies or by direct fluorescence using an Odyssey Clx imaging system (LI-COR) after incubation with secondary antibodies coupled to a fluorescent dye (Supplementary Table 2).

### Colony formation assay

A549 cells were plated on 6 well plates with 1 × 10^5^ cells per well and treated with the respective damaging agents for 1 h. 24 h later, cells were re-plated on 10 cm dishes with 300 cells per dish and grown for 14 days. Colonies were stained with a 0,1% crystal violet solution and counted manually. Cell survival was normalized to the colony count of untreated cells.

### Irradiation of cells

A cabinet X-ray irradiator (CellRad instrument, Faxitron Bioptics LLC) was used with an automated dose control setting at 130 kV and 5 mA to deliver a preset total X-ray dose.

### Laser micro-irradiation followed by live-cell imaging

For laser micro-irradiation microscopy, U2OS cells were seeded in 35 mm dishes (Ibidi) and transfected with the respective GFP-Myo6 expression plasmids using Lipofectamine 2000 (Life technologies) according to the manufactureŕs instructions. An mCherry-XRCC4 expression plasmid was co-transfected and served as control for successful induction of double strand breaks. Laser micro-irradiation was carried out on a Visitron Spining Disk microscope equipped with an environmental chamber set to 37°C. DSB-containing tracks were generated using 2% power (corresponding to 6nW average power measured at the microscope back port) of a 355 nm pulsed laser (STV-01E-1×0, Teem Photonics). Images were recorded with a 60x water immersion objective (CFI Plan Apo VC 60XWI, Nikon) for 5 min following irradiation at intervals of 5 s. A custom macro in Fiji/ImageJ was used to measure the accumulation of fluorescently tagged Myo6 variants at micro-irradiation sites from cells that showed XRCC4 recruitment to the damaged area. The recruitment was calculated as the ratio of the mean fluorescent intensities of the ROI over the nuclear signal.

### Quantification of Laser Micro-Irradiation Data

Time-lapse fluorescence microscopy data were analyzed using a custom Fiji/ImageJ macro designed to automate processing of laser micro-irradiation experiments. The macro processes image stacks and corresponding ROI files, identifying irradiated regions. It detects the first and last frames of micro-irradiation by identifying significant intensity changes and removes frames with signal from the laser. Intensity profiles are extracted along ROIs, which are expanded for averaging based on user-defined widths. Background subtraction is performed using a narrow ROI centered on the signal and a wider ROI encompassing both signal and background. Here, we used a ROI line width of 16 pixels for averaging of the signal, and ROI widths for background estimation of 18 and 24 pixels. The background intensity is calculated as the mean of the wider ROI excluding the narrower one.

Curves that showed fluctuations in the background signal (e.g. bright nucleoli entering the ROIs, cell movement) were removed from the analysis. We calculated the log-fold change (logFC) of the fluorescence signal at the irradiated region relative to the nuclear background signal. The time point t = 0 corresponds to the first frame after irradiation. For statistical analysis, we fitted a non-linear mixed effects regression model using the R packages *nlme*^34,35^ and *emmeans*^36^. The model equation describes exponential rise to steady state, implemented with the self-starting asymptotic model *SSasymp()* in R:

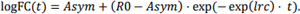

Here, the parameter *Asym* denotes the asymptote, *R0* the initial value, and *lrc* the natural logarithm of the rate coefficient (constraining the rate to be positive). The nonlinear mixed effects model included three predictive factors: *genotype*, *replicate*, and individual *curve*. The primary factor of interest, *genotype*, was included as a fixed effect. To avoid pseudo-replication and account for unwanted sources of variability, the nested factors *replicate* and *curve* were included as random effects. We assumed a diagonal covariance structure for both random effects, implying independent effects on the model parameters. To account for temporal autocorrelation, a continuous autoregressive correlation structure of order 1 (corCAR1) was applied. Model parameter estimates were compared between genotypes using pairwise contrasts. P-values were adjusted for multiple testing using a multivariate t correction (“mvt” in *emmeans*). Model selection was guided by the Akaike Information Criterion (AIC) and likelihood ratio tests.

### Traffic Light Reporter (TLR) assay

The TLR assay was performed in U2OS cells as described by Certo et al^23^. pCVL Traffic Light Reporter 1.1 (Sce target) Ef1a Puro plasmid (TLR) (Addgene Cat# 31482) was integrated into U2OS cells by lentiviral transduction^23^. TLR-integrated U2OS cells were selected with 2 µg/ml puromycin for 2 weeks. U2OS-TLR cells were plated on 10 cm dishes and transfected using Lipofectamine 2000 according to the manufacureŕs instructions to deliver SceI (pRRL sEF1a HA.NLS.Sce(opt). T2A.IFP, Addgene #31484), the Donor template (pRRL SFFV d20GFP.T2A.mTagBFP Donor, Addgene #31485) and respective siRNAs for depletion of Myo6 and CtIP (Supplementary Table 3). Cells were harvested 72 h post-transduction and analyzed on a BD LSRFortessa SORP flow cytometer. To identify SceI nuclease and GFP-repair-template-positive cells, IFP1.4 was measured using a 640 nm laser and a 730/45 filter, and mTagBFP was measured using a 405 nm laser and a 450/50 filter. Cells positive for IFP and BFP were further analyzed for GFP (488 nm laser, 530/30 filter) and mCherry (561 nm laser, 610/20 filter) expression. To determine the corresponding cell cycle stage, half of the harvested cells were fixed in ice-cold ethanol, stained with Propidium Iodide, and analyzed by flow cytometry using a 561 nm laser and a 610/20 filter. Data were analyzed using FlowJo software (v10.10, BD). Knockdown efficiency was monitored by western blotting using specific antibodies against CtIP and Myo6 (Supplementary Table 2).

### DSB mobility

U2OS cells expressing Rad52 fused to mCherry were seeded into a 35 mm glass-bottom dishes (MatTek) in DMEM supplement with 10%FBS, 100 U/mL penicillin, 100 µg/mL streptomycin and 200 mM GlutaMAX. Medium was changed to FluoroBrite DMEM supplemented with 5 % FBS, 100 U/mL penicillin, 100 µg/mL streptomycin, 200 mM GlutaMAX and 400 mM HEPES 1 h before the start of imaging. Cells were imaged 20 h after X-ray irradiation (dose rate of 1 Gy/26 s, 250 keV, 10 mA) on a Nikon Eclipse Ti-E inverted microscope using a 60x plan apochromat objective and a DS-Qi2 camera. Images comprising 5 z-stacks with a step size of 0.3 µm were taken every 5 min over a period of 90 min using Nikon NIS Elements AR software. Fluorescent proteins were excited with a X-Cite® 120 LED illumination system (Lumen Dynamics) using the following filter sets: 562/40 nm excitation, 593 nm beam splitter and 624/40 emission filter. The microscope was enclosed with an incubation chamber with constant temperature (37°C), atmosphere (CO_2_ concentration 5 %), and humidity (Okolab). Image analysis was performed on maximum intensity projections of z-stacks preprocessed by flat field correction. Individual nuclei were select in Fiji and registration was performed using the plugin StackReg. Foci were followed using TrackMate^37^. Tracks with a minimum length of 5 time points were used to calculate mean square displacements (MSD) and diffusion coefficients using the MSDanalyzer class in Matlab^38^.

### Gene Ontology (GO) term enrichment analysis

Proteins identified in the MyUb SILAC interactome experiment were filtered to select enriched interactors (fold change > 2, FDR < 0.05, detected in more than one replicate). Using R, gene symbols were extracted from UniProt annotations and mapped to Entrez Gene IDs using the org.Hs.eg.db package^39^. GO enrichment for biological processes was performed with clusterProfiler using custom gene sets defined for “homologous recombination” (HR, GO:0000724) and “non-homologous end joining” (NHEJ, GO:0006303), with all annotated human genes as background. Pathway assignments were extracted from GO-parent relationships via org.Hs.eg.db, and enrichment statistics were calculated with the enricher function, using Benjamini–Hochberg correction for multiple testing. Bar plots of enriched pathway members, as well as the volcano plot, were generated with ggplot2.

### Protein production and purification

GST fusion proteins were produced in *E. coli* Bl21 (DE3) cells. Expression was induced with 1 mM IPTG (Generon) at an OD_600_ of 0.8 at 37°C for 4 h. Cells were pelleted and lysed by sonication in PBS/0.1% Triton X-100 (Merck) supplemented with protease inhibitor cocktail (SIGMAFAST). Clarified supernatants were incubated with 1 ml of GSH-Sepharose beads (Cytiva) per liter of bacterial culture. After 2 h at 4°C, the beads were washed with PBS/0.1% Triton X-100 and re-buffered in storage buffer (50 mM Tris, pH 7.4, 100 mM NaCl, 1 mM EDTA, 1 mM DTT, and 10% glycerol). VCP was purified as GST-3C-fusion followed by cleavage of the GST-tag by ^GST^PreScission protease and size-exclusion chromatography (SEC).

Co-expression of His-KU80 with untagged KU70 was performed in *E. coli* BL21 (DE3) pLysS at 16°C for 20 h after induction with 1 mM IPTG. Cells were resuspended in buffer A (50 mM Tris-HCl pH 7.4, 250 mM NaCl, 10% glycerol, 1 mM DTT, 20 mM imidazole) and lysed by sonication. The clarified supernatant was subjected to affinity chromatography on Ni-NTA resin (Qiagen), and eluted protein was rebuffered using PD 10 columns (Cytiva) in JS buffer (100 mM HEPES pH 7.5, 50 mM NaCl, 5% glycerol, 1% Triton X-100, 2 mM MgCl_2_, 5 mM EGTA, 1 mM DTT).

### Preparation of unanchored K63-linked polyubiquitin chains

Unanchored K63-linked polyubiquitin chains were prepared as previously described^40^. Briefly, chains were produced in a 1 ml reaction containing 40 mM HEPES, pH 7.4, 8 mM magnesium acetate, 50 mM NaCl with 30 µM ATP, 0.05 µM E1 (^His^Uba1), 2 µM ^His^Ubc13-Mms2 and 0.5 µM E3 (Pib1RING+100aa). *Wildtype* ubiquitin (purified bovine ubiquitin, Sigma) was used at a concentration of 8 µM, and 4 µM of ubiquitin mutant K63R was added for capping of the chains. The reaction was incubated for 1.5 h at 30°C, and 1-4 µl of the reaction mix were used in GST-pulldown assays.

### GST-pulldown assays

GST-pulldown assays were performed by incubating 10 µg GST (as control) or 14 µg GST-MyUb immobilized on 20 µl GSH-Sepharose beads with lysates from 5×10^6^ unlabeled HeLa cells, and interactors were detected by western blotting using antibodies against endogenous proteins.

To identify direct protein-protein interactions, we performed pulldown assays by incubating 5 µg GST or GST-UBDs (MyUb or MIUMyUb) immobilized on 20 µl GSH-Sepharose beads with various concentrations of either His-KU70/80 complex, K63-linked ubiquitin chains or VCP either alone or in combinations in 200 µl modified JS buffer (100 mM HEPES pH 7.5, 50 mM NaCl, 5% glycerol, 1% Triton X-100, 2 mM MgCl_2_, 5 mM EGTA, 1 mM DTT). Beads were washed 3 times in 1 ml modified JS-buffer, boiled for 10 min in NuPAGE^®^ LDS Sample Buffer and subjected to SDS-PAGE and subjected to western blotting.

### AlphaFold 3 predictions

The AlphaFold 3 Server^33^ was queried with the MyUb domain of Myo6 in complex with designed fragments of KU70, KU80 and VCP. The fragments were designed based on TED-domain annotation^41^ (from the AlphaFold Database (https://pubmed.ncbi.nlm.nih.gov/37933859/). Per Fragment of either VCP, KU70 or KU80 one run was performed initially in complex with MyUb with a random seed and default template settings. Likelihood of interaction between MyUb and each respective fragment from KU70, KU80 and VCP was evaluated by means of the predicted alignment error (pAE) between the MyUb domain and each respective query. For each query with low pAE between MyUb and the query fragment and an iPTM in the range of 0.7 a second prediction was performed with the same settings and a new randomly generated seed.

## Acknowledgements

We thank Prof. Markus Löbrich for scientific discussions. We thank Les Hanakahi, Simona Polo, Martijn Luijsterburg and Christian Renz for sharing constructs, antibodies, cell lines or recombinant proteins. The IMB Core Facilities for Proteomics, Flow Cytometry, Microscopy and Protein Production are acknowledged for technical support and reagents. Funding of the German Research Foundation supported the lnvitrogen Bigfoot Cell Sorter (P#511658729, IMB Flow Cytometry Core Facility), the BD LSRFortessa SORP (P#210253511, IMB Flow Cytometry Core Facility), the AF7000 widefield microscope (P#212049334, IMB Microscopy Core Facility) and the Visitron Spinning Disk microscope (P#402386039, IMB Microscopy Core Facility).

This work was funded by the Deutsche Forschungsgemeinschaft (DFG, German Research Foundation)—Project-ID 393547839—SFB 1361 awarded to H.D.U., A.L., V.R. and P.B., CRC1292 (TP05) awarded to K.R., Project-ID BE 5342/2-1—FOR 2800 awarded to P.B. and Project-IDs 408799149 and 555596903 awarded to H.P.W.

## Author contributions

H.D.U., V.R. and H.P.W. conceived the study, K.H., J.S., R.P.W., F.F.S., S.R., A.A.K., L.M.V., M.S.S., T.S., S.N. and H.P.W. performed experiments and data analysis in cell and molecular biology, I.M. and P.B. performed mass spectrometry data analysis, A.L. and S.R. designed and performed DSB mobility experiments and data-analysis, M.G. provided technical support and wrote the quantification script for laser-irradiation experiments, F.K. performed their statistical analyses. K.R. provided resources. H.D.U. and H.P.W. wrote the manuscript and created the figures, and all authors discussed the data and provided input during manuscript preparation.

## Data availability

All reagents used in the paper are listed in Supplementary Table 5.

## Code availability

Custom codes used for the Image J-based quantifications of RPA32, RAD51, BrdU and KU80 foci and the quantification of laser-irradiation images are available on GitHub (https://github.com/hwollsch/Myo6-in-DSB-repair). The code for MSD analysis and the respective data is accessible via the institutional repository of Technical University Darmstadt (https://doi.org/10.48328/tudatalib-1981).

