## Supplementary material for "The nuclear actin cytoskeleton supports DNA double-strand break repair via VCP-mediated extraction of the KU70/80 complex from damaged chromatin": Supplementary_material_Hauschulte_etal.pdf

[#] Equal contribution

1. Supplementary figures and legends

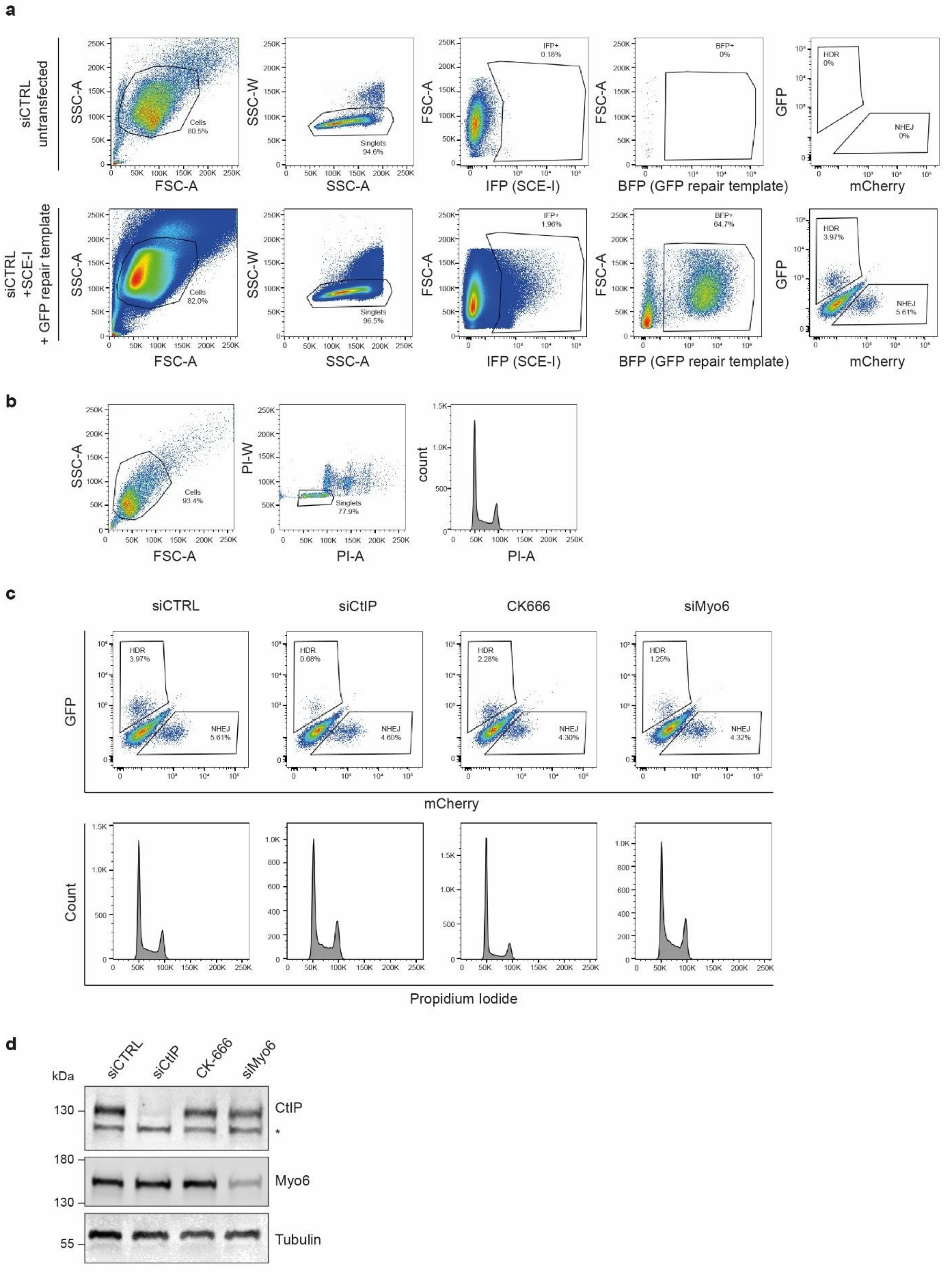


**Fig. S1. The Traffic Light Reporter (TLR) assay reveals a function of Myo6 in HDR**

**a,** Representative gating for the TLR assay shown in **Fig. 1c**. Cells were sequentially gated for the cell population (FSC-A vs SSC-A), singlets (SSC-W vs SSC-A), SCE-I transfected cells (IFP$^+$), and GFP repair template-positive cells (BFP$^+$). Homologous recombination (HR, GFP$^+$) and non-homologous end joining (NHEJ, mCherry$^+$) outcomes were quantified. Data shown for untransfected siCTRL cells serve as gating cutoff.

**b,** Gating for cell cycle analysis for the TLR assay shown in **Fig. 1c**. Cells were gated by FSC-A vs SSC-A, singlets were selected (PI-W vs PI-A), and DNA content was assessed by propidium iodide (PI) staining.

**c,** Representative results of TLR assay. Cells were treated with control siRNA (siCTRL), CtIP knockdown (siCtIP), Arp2/3 inhibitor CK666, or Myosin VI knockdown (siMyo6). Flow cytometry plots show relative HR and NHEJ frequencies. Corresponding cell cycle profiles, assessed by propidium iodide staining, are displayed below.

**d,** Western blots to monitor knockdown efficiencies for the TLR assay shown in **Fig1c**. U-2 OS TLR cells were either treated with 50 µM of the ARP2/3 inhibitor CK666 for 72 h or co-transfected with siRNAs against CtIP or Myo6 as indicated and analyzed by western blotting with the indicated antibodies 72 h post-transfection. The asterisk indicates unspecific binding of the CtIP antibody.

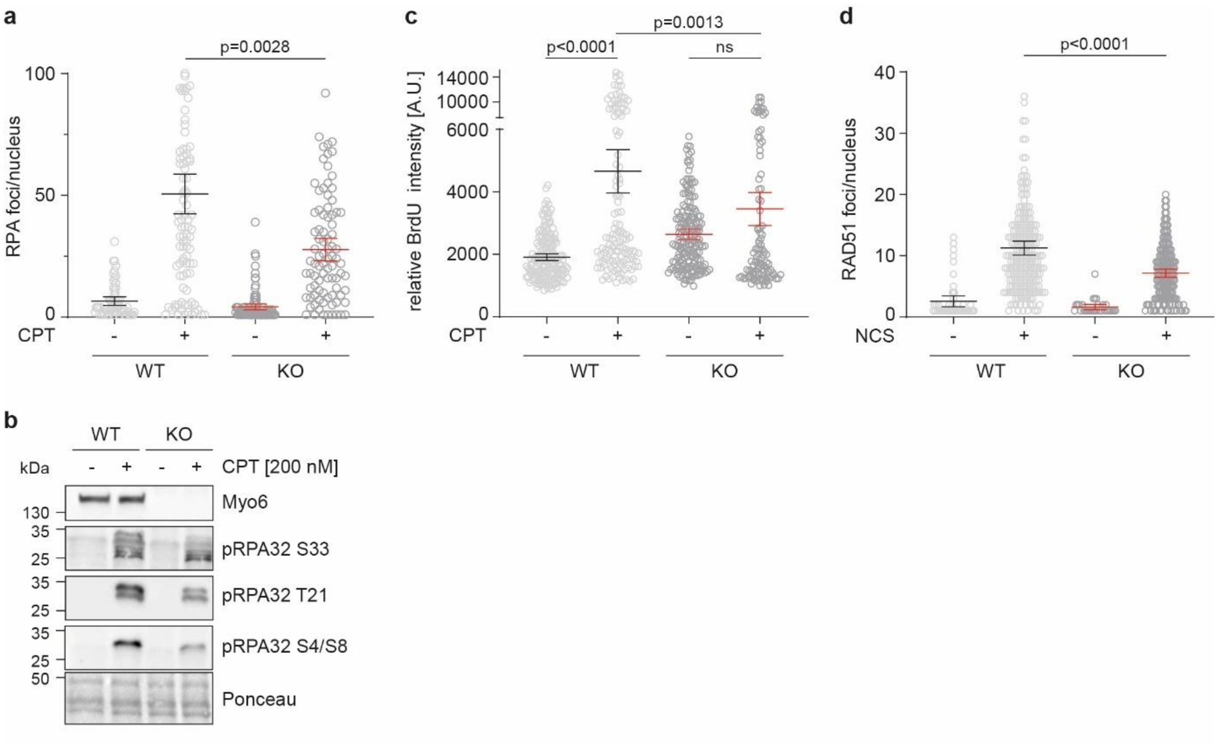


**Fig. S2. Myo6 promotes efficient resection of DSBs**

**a,** Myo6 promotes RPA foci formation upon DSB induction. A549 WT and Myo6 KO cells were treated with 1 µM CPT for 1 h, followed by immunofluorescence microscopy against RPA32.

**b,** Myo6 promotes RPA32 phosphorylation upon DSB induction. A549 WT and Myo6 KO cells were treated with 200 nM CPT for 1 h. Total cell lysates were analyzed by western blotting and Ponceau S staining.

**c,** Myo6 promotes ssDNA formation upon DSB induction. A549 WT and Myo6 KO cells were labeled with BrdU for 24 h, treated with 1 µM CPT for 1 h and analyzed by immunofluorescence microscopy under non-denaturing conditions using a BrdU-specific antibody.

**d,** Myo6 promotes RAD51 foci formation upon DSB induction. A549 WT and Myo6 KO cells were treated with 1 µg/ml NCS for 1 h followed by immunofluorescence microscopy against RAD51.

For **a-d,** A representative experiment from at least three independent replicates is shown.

For **a, c, d,** Signal intensities were quantified using Image J and shown as dot plots with mean values −/+ 95% confidence intervals. Significance levels were calculated using the Mann-Whitney test (ns: not significant).

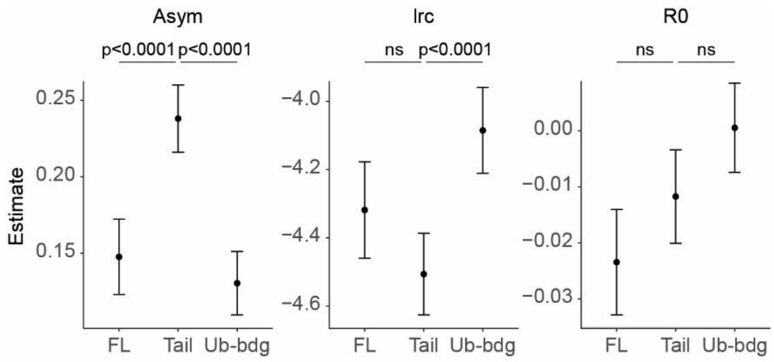


**Fig. S3. Summary of parameter estimates from the nonlinear mixed-effects model**

*Asym* is the model asymptote, *R0* the initial value, and *lrc* the logarithm of the rate coefficient. Quantitatively, *Asym* is the log-fold change of the signal at the irradiated region relative to nuclear background at steady-state. Error bars indicate 95%-confidence intervals. P-values are from pairwise contrasts, adjusted for multiple testing across all genotype comparisons.

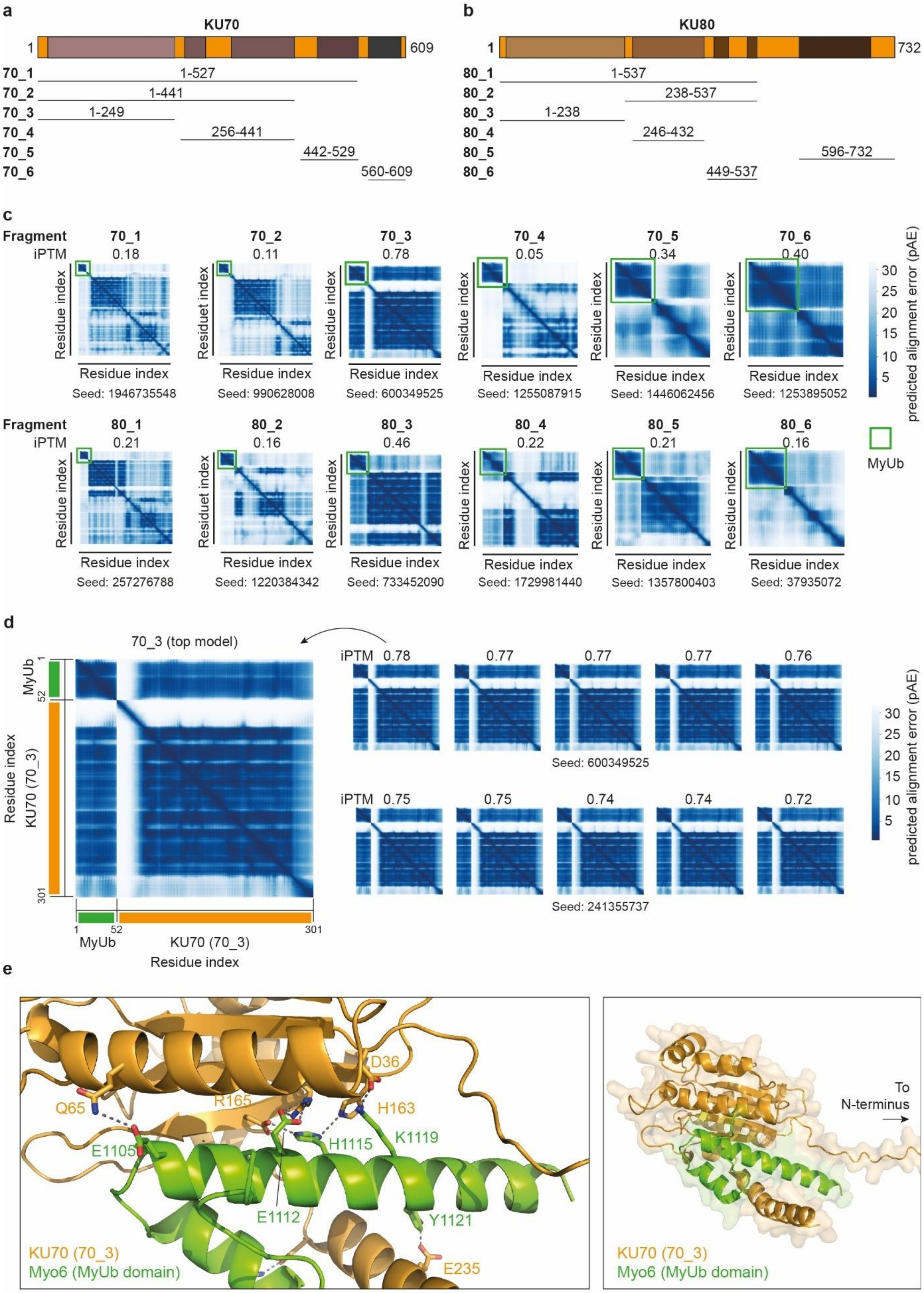


**Fig. S4. AlphaFold 3 predicts an interaction of Myo6 with the N-terminus of KU70**

**a,** Schematic representation of K70 fragments used for MyUb interaction screening.

**b,** Schematic representation of KU80 fragments used for MyUb interaction screening.

**c,** AlphaFold 3 predicts an efficient interaction of MyUb only with one N-terminal fragment of KU70 (70-3, aa 1-249). The AlphaFold3 server was queried with MyUb and each KU70 and KU80 fragment using randomly generated seeds, and iPTM scores and pAE matrices are plotted for the top model of each run.

**d,** AlphaFold 3 models of MyUb in complex with KU70 fragment 70_3 show good agreement. Plotted are pAE matrices and iPTM scores for ten models, derived from two queries using different seeds.

**e,** Predicted interface between KU70 (fragment 70_3, orange) and MyUb (green). Residues involved in the interaction and possible hydrogen bonds and salt bridges are labeled. MyUb is pinched between the main body of the N-terminal folded domain of KU70 and one additional helix. The interface is predicted to be stabilized by several electrostatic interactions, including two salt bridges.

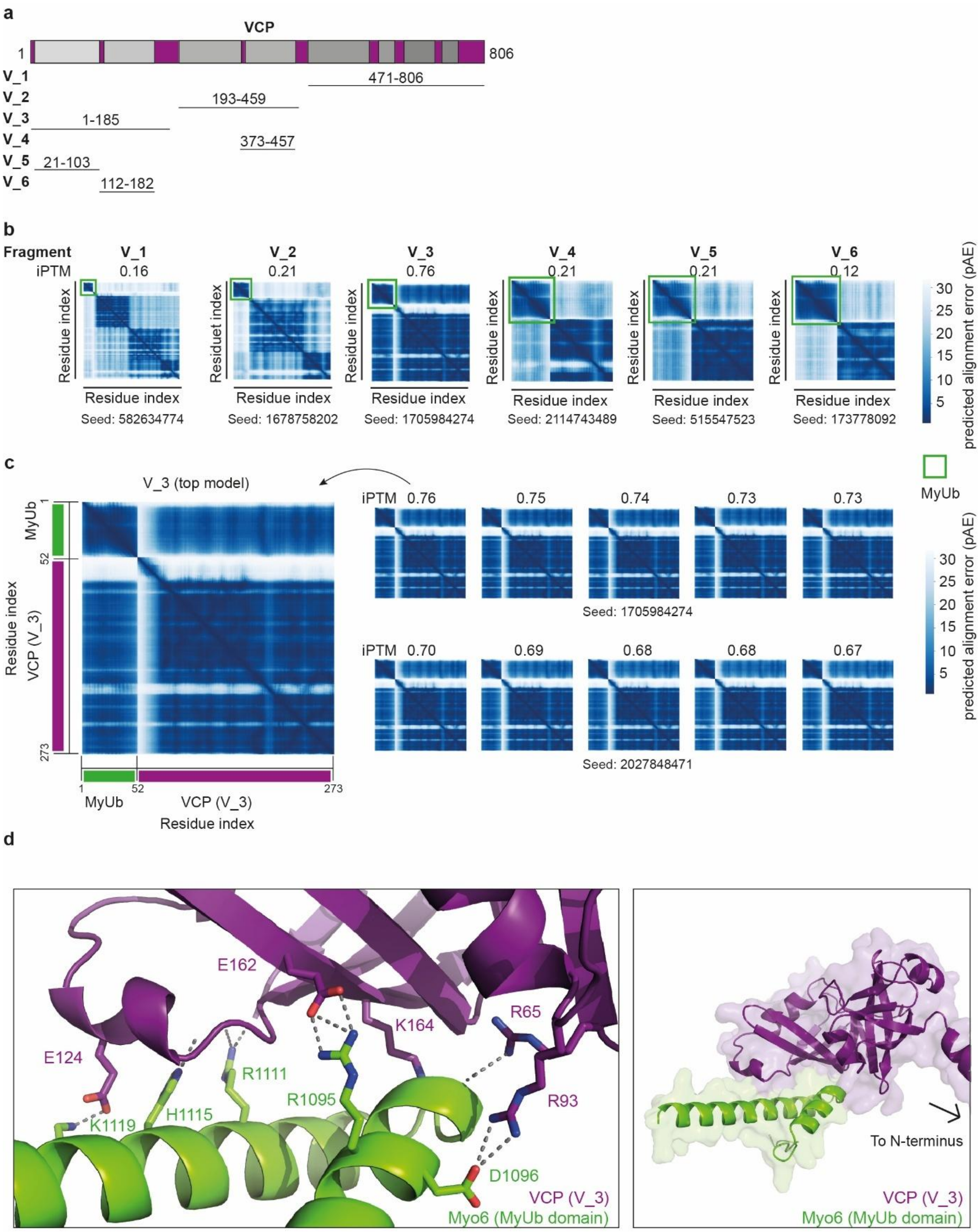


**Fig. S5. AlphaFold 3 predicts an interaction of Myo6 with the N-terminus of VCP**

**a,** Schematic representation of VCP fragments used for MyUb interaction screening.

**b,** AlphaFold 3 predicts an efficient interaction of MyUb only with one N-terminal fragment of VCP (V-3, aa 1-185). The AlphaFold3 server was queried with MyUb and each VCP fragment using randomly generated seeds, and iPTM scores and pAE matrices are plotted for the top model of each run.

**c,** AlphaFold 3 models of MyUb in complex with VCP fragment V_3 show good agreement. Plotted are pAE matrices and iPTM scores for ten models, derived from two queries using different seeds.

**e,** Predicted interface between VCP (fragment V_3, purple) and MyUb (green). Residues involved in the interaction and possible hydrogen bonds and salt bridges are labeled. The interface is predicted to be stabilized by several electrostatic interactions, including two salt bridges.

### 2. Supplementary Tables

**Supplementary Table 1. Plasmids used in this study**

| Plasmid | Source | Identifier |
|---|---|---|
| pGEX6P1 | Merck | Cytiva 28-9546-48 |
| pGEX6P1-MyUb | He et al.[1] | 2893 |
| pGEX6P1-MIUMyUb | Wollscheid et al.[2] | n/a |
| pEGFP-C1 | Clontech | 2535 |
| pOG44 –Flp-Recombinase | Thermo Fisher Scientific | V600520 |
| pEGFP-C1-myo6 (fl) | Wollscheid et al.[2] | 2905 |
| pEGFP-C1-myo6-tail (aa732-1262) | Wollscheid et al.[2] | 2908 |
| pEGFP-C1-myo6-tail ubiquitin binding mutant (A1013G+I1072A*) | Shi et al.[3] | 5358 |
| pET3a-Ubi(K63R) | Parker et al.[4] | 684 |
| pET21(+)-His6-Ubc13 | Renz et al.[5] | 3010 |
| pET30-His6-TEV-Mms2 | Renz et al.[5] | 2807 |
| pET30-His6-TEV-Pib1RING+100aa | Renz et al.[5] | 3019 |
| pET28-mE1 (ubiquitin activating enzyme E1) | Carvalho et al.[6] | 2448 |
| XRCC4-mCherry | Luijsterburg lab | 5957 |
| pEGFPC1-VCP | This study | 3108 |
| pGex6P1-VCP | This study | 3387 |
| KU70/80 in HispET | Les Hanakahi[7] | 5707 |
| pCVL Traffic Light Reporter 1.1 (Sce target) Ef1a Puro | Certo et al.[8], Addgene #31482 | 3498 |
| pRRL sEF1a HA.NLS.Sce(opt).T2A.IFP | Certo et al.[8], Addgene #31484 | 3499 |
| pRRL SFFV d20GFP.T2A.mTagBFP Donor | Certo et al.[8], Addgene #31484 | 3500 |
| pEGFP-C1-FLAG-Ku70 | Britton et al[9]. Addgene #46957 | 5661 |
| pEGFP-C1-FLAG-Ku80 | Britton et al[9]. Addgene #46958 | 5662 |
| pEGFPC1_KU70_1-248 | This study | 6582 |
| pEGFPC1_VCP1-104 | This study | 6551 |

*I1072 in this construct (isoform 2) corresponds to I1104 in the long isoform (3). Mutation to alanine abrogates ubiquitin binding of the MyUb domain.

**Supplementary Table 2. Antibodies used in this study**

| Antibody | Source | Identifier |
|---|---|---|
| Mouse monoclonal anti-GFP (clone 7.1/13.1) | Roche | Cat#11814460001; RRID: AB_390913 |
| Mouse monoclonal anti-Myosin VI (clone MUD19) | Merck | Cat#M0691; RRID:AB_369989 |
| Rabbit polyclonal anti-Myosin VI | Wollscheid et al.[2] | n/a |
| Rabbit polyclonal anti-CtIP | Bethyl Laboratories | Cat#A300-488A-M RRID:AB_2175262 |
| Mouse monoclonal anti-RPA2 (clone 9H8) | Invitrogen | Cat#MA1-26418; RRID:AB_795362 |
| Rabbit polyclonal anti-Phospho-RPA2 T21 | R and D Systems (Biotechne) | Cat#AF6654; RRID:AB_10888653 |
| Rabbit polyclonal anti-Phospho-RPA32 S4/S8 | Bethyl Laboratories | Cat# A300-245A; RRID:AB_210547 |
| Rabbit polyclonal anti-Phospho-pRPA S33 | Bethyl Laboratories | Cat# A300-246A-M; RRID:AB_210547 |
| Rabbit polyclonal anti-VCP (clone 7F3) | Cell Signaling Technology | Cat#2649; RRID:AB_2214629 |
| phospho-Histone H2A.X (Ser139) gammaH2AX (clone JBW301) | Millipore | Cat# 05-636-I; RRID |
| Mouse monoclonal anti-BrdU (clone B44) (IdU) | BD Biosciences | Cat#347580; RRID:AB_400326 |
| Rabbit monoclonal anti-Rad51 (clone D4B10) | Cell Signaling Technology | Cat#8875; RRID:AB_2721109 |
| KU80 | | |
| KU70 | | |
| Ubiquitin (P4D1) | Cell Signaling Technology | Cat#3936; RRID:AB_331292 |
| Rabbit polyclonal anti-phospho-Chk1 (Ser345) | Cell Signaling Technology | Cat#2341; RRID:AB_330023 |
| Tubulin | | |
| IRDye® 680LT donkey anti-rabbit IgG secondary antibody | LICOR | Cat#926-68023; RRID: AB_10706167 |
| IRDye® 680LT donkey anti-mouse IgG secondary antibody | LICOR | Cat#926-68072; RRID: AB_10953628 |
| IRDye® 800CW goat anti-rabbit IgG secondary antibody | LICOR | Cat#926-32211; RRID: AB_621843 |
| IRDye® 800CW donkey anti-mouse IgG secondary antibody | LICOR | Cat#926-32212; RRID: AB_621847 |
| Polyclonal goat anti-goat HRP secondary antibody | Dako | Cat# P044901-2; RRID:AB_2617143 |
| Polyclonal goat anti-mouse HRP secondary antibody | Dako | Cat#P044701-2; RRID:AB_2617137 |
| Polyclonal goat anti-rabbit HRP secondary antibody | Dako | Cat#P044801-2; RRID:AB_2617138 |
| Goat anti-Rat IgG (H+L) Secondary Antibody, Alexa Fluor 488 | Thermo Fisher Scientific | Cat#A-11006; RRID:AB_2534074 |
| Goat anti-Mouse IgG (H+L) Secondary Antibody, Alexa Fluor 647 | Thermo Fisher Scientific | Cat#A-21236; RRID:AB_2535805 |

| | | |
|---|---|---|
| Goat anti-Rabbit IgG (H+L) Cross-Adsorbed, Alexa Fluor 647 | Thermo Fisher Scientific | Cat#A-21244; RRID:AB_2535812 |

**Supplementary Table 3. siRNAs used in this study**

| siRNA | Source | Identifier/Sequence |
|---|---|---|
| Allstars Negative control siRNA | Qiagen | Cat#SI03650318 |
| Hs_MYO6_5 FlexiTube siRNA | Qiagen | Cat#SI03142692 |
| Hs_MYO6_7 FlexiTube siRNA | Qiagen | Cat#SI04243351 |
| Hs_MYO6_8 FlexiTube siRNA | Qiagen | Cat#SI04370737 |
| Hs_MYO6_10 FlexiTube siRNA | Qiagen | Cat#SI04998749 |
| CtIP | Dharmacon | GCUAAAACAGGAACGAAUCUU |

**Supplementary Table 4. Oligonucleotides used in this study**

| Oligonucleotides for mutagenesis | Sequence (5´- 3´) | Construct | ID |
|---|---|---|---|
| KU70 E250 STOP For | GCGGAAGGTTCGCGCCAAGTAAACAAGGAAGCGAGCACTCAGC | pEGFP-C1-KU70 E250 → STOP | 7935 |
| KU70 E250 STOP Rev | GCTGAGTGCTCGCTTCCTTGTTTACTTGGCGCGAACCTTCCGC | | 7936 |
| VCP P104 STOP For | GGGGATGTCATCAGCATCCAGTAATGCCCTGATGTGAAGTACGGC | pEGFP-C1-VCP P104 → STOP | 7941 |
| VCP P104 STOP Rev | GCCGTACTTCACATCAGGGCATTACTGGATGCTGATGACATCCCC | | 7942 |
| **Oligonucleotide for amplification** | **Sequence (5´- 3´)** | **Construct** | **ID** |
| VCP_For_BglII | GAAGATCTATGGCTTCTGGAGCCGATTCAAAAGG | pEGFP-C1-VCP and pGex6P1-VCP | 3733 |
| VCP_Rev_XbaI_BamHI | GCTCTAGAGGATCCTTAGCCATACAGGTCATCATCATTGTCTTCTGTG | | 3734 |

**Supplementary Table 5. Reagents used in this study**

| Chemical | Source | Identifier |
|---|---|---|
| CK666 | Merck | SML0006-25MG |
| Camptothecin | Merck | C9911-100MG |
| Neocarcinostatin | Merck | N9162-100UG |
| NMS-873 | Merck | SML1128-5MG |
| IPTG | Generon | Cat#GEN-S-02122 |
| Ni-NTA agarose | Qiagen | Cat#30250 |
| Imidazole | Merck | Cat#I2399 |
| Glutathione | Merck | Cat#G4251 |
| Paraformaldehyde | Merck | Cat#P6148 |
| Formaldehyde solution, 36.5-38% in H2O | Merck | Cat#F8775 |
| SIGMAFAST protease inhibitor cocktail | Merck | Cat#S8830 |
| cOmplete Protease Inhibitor Cocktail | Roche | Cat#5056489001 |
| Pfu Turbo DNA Polymerase | Agilent | Cat#600250 |
| DpnI | New England Biolabs | Cat#R0176L |
| Lipofectamine® 2000 | Thermo Fisher Scientific | Cat#11668019 |
| Lipofectamine® RNAiMAX | Thermo Fisher Scientific | Cat#13778150 |
| FuGENE® HD Transfection Reagent | Promega | Cat#E2311 |
| ProLong™ Diamond Antifade Mountant | Thermo Fisher Scientific | Cat#15205739 |
| DMEM, high glucose, pyruvate, no glutamine | Thermo Fisher Scientific | Cat#21969035 |
| Trypsin-EDTA (0.05%), phenol red | Thermo Fisher Scientific | Cat#25300054 |
| Penicillin-Streptomycin (10,000 U/mL) | Thermo Fisher Scientific | Cat#15140122 |
| L-Glutamine (200 mM) | Thermo Fisher Scientific | |
| Triton X-100 | Merck | Cat#T9284 |
| Albumin (bovine serum albumin, BSA) | Merck | Cat#A7906 |
| Sodium deoxycholate | Merck | Cat#D6750 |
| SDS, 20%, Sodium dodecyl sulfate | Merck | Cat#05030 |
| NuPAGE LDS Sample Buffer (4X) | Thermo Fisher Scientific | Cat#11559166 |
| GFP-Trap magnetic agarose beads | Chromotek | Cat#gtma-100 |
| Milk powder, skim milk | Merck | Cat#70166 |
| Tween-20 | Merck | Cat#P7949 |
| Amersham ECL Select Western Blotting Detection Reagent | Thermo Fisher Scientific | Cat#RPN2235 |
| Amersham ECL Prime Western Blotting Detection Reagent | Thermo Fisher Scientific | Cat#12994780 |
| Ponceau S | Merck | Cat#P3504 |
| Hoechst 33342 | Thermo Fisher Scientific | Cat#11534886 |
| PageRuler Prestained Protein Ladder | Thermo Fisher Scientific | Cat#11822124 |
| 4-15% Criterion™ TGX Stain-Free™ Protein Gel, 26 well, 15 µl | Bio-Rad Laboratories | Cat#567-8085 |
| Mini-PROTEAN TGX Stain Free Gels, 4-15%, 15-well | Bio-Rad Laboratories | Cat#456-8086 |
| Glutathione Sepharose High Performance resin (GSH) | Cytiva | Cat#17527901 |
| Recombinant ubiquitin activating enzyme (E1) | In house protein production core facility | Carvalho et al[6] |
| Ubiquitin from bovine erythrocytes | Merck | Cat#U6253-25MG |
| 35mm dishes | Ibidi | Cat#80136 |
| **Critical commercial assays** | | |

| | | |
|---|---|---|
| Trans-Blot® Turbo™ RTA Midi Nitrocellulose Transfer Kit | Bio Rad laboratories | Cat#1704271 |
| Gateway® LR Clonase® II enzyme mix | Thermo Fisher Scientific | Cat#10134992 |
